# Structural and Mechanistic Bases for Resistance of the M66I Capsid Variant to Lenacapavir

**DOI:** 10.1101/2024.11.25.625199

**Authors:** Lorenzo Briganti, Arun S. Annamalai, Stephanie M. Bester, Guochao Wei, Jonathan R. Andino-Moncada, Satya P. Singh, Alex B. Kleinpeter, Meghna Tripathi, Binh Nguyen, Rajalingam Radhakrishnan, Parmit K. Singh, Juliet Greenwood, Lauren I. Schope, Reed Haney, Szu-Wei Huang, Eric O. Freed, Alan N. Engelman, Ashwanth C. Francis, Mamuka Kvaratskhelia

**Author notes:** Corresponding author: Mamuka Kvaratskhelia. These authors contributed equally to this work (Lorenzo Briganti, Arun S. Annamalai). Current address: Department of Microbiology & Infectious Disease Center, School of Basic Medical Sciences, Peking University, Beijing, China (Guochao Wei). Current address: Emory University School of Medicine, GA, USA. Current address: Nuffield Department of Medicine, University of Oxford, Oxford, UK. Current address: University of Southern California, CA, USA.

## Abstract

Lenacapavir (LEN) is the first in class viral capsid protein (CA) targeting antiretroviral for treating multi-drug-resistant HIV-1 infection. Clinical trials and cell culture experiments have identified resistance associated mutations (RAMs) in the vicinity of the hydrophobic CA pocket targeted by LEN. The M66I substitution conferred by far the highest level of resistance to the inhibitor compared to other RAMs. Here we investigated structural and mechanistic bases for how the M66I change affects LEN binding to CA and viral replication. The high-resolution X-ray structure of the CA(M66I) hexamer revealed that the β-branched side chain of Ile66 induces steric hindrance specifically to LEN thereby markedly reducing the inhibitor binding affinity. By contrast, the M66I substitution did not affect binding of Phe-Gly (FG)-motif-containing cellular cofactors CPSF6, NUP153, or SEC24C, which engage the same hydrophobic pocket of CA. In cell culture the M66I variant did not acquire compensatory mutations or replicate in the presence of LEN. Analysis of viral replication intermediates revealed that HIV-1_(M66I CA)_ predominantly formed correctly matured viral cores, which were more stable than their wildtype counterparts. The mutant cores stably bound to the nuclear envelope but failed to penetrate inside the nucleus. Furthermore, the M66I substitution markedly altered HIV-1 integration targeting. Taken together, our findings elucidate mechanistic insights for how the M66I change confers remarkable resistance to LEN and affects HIV-1 replication. Moreover, our structural findings provide powerful means for future medicinal chemistry efforts to rationally develop second generation inhibitors with a higher barrier to resistance.

**IMPORTANCE:** Lenacapavir (LEN) is a highly potent and long-acting antiretroviral that works by a unique mechanism of targeting the viral capsid protein. The inhibitor is used in combination with other antiretrovirals to treat multi-drug-resistant HIV-1 infection in heavily treatment-experienced adults. Furthermore, LEN is in clinical trials for preexposure prophylaxis (PrEP) with interim results indicating 100 % efficacy to prevent HIV-1 infections. However, one notable shortcoming is a relatively low barrier of viral resistance to LEN. Clinical trials and cell culture experiments identified emergent resistance mutations near the inhibitor binding site on capsid. The M66I variant was the most prevalent capsid substitution identified in patients receiving LEN to treat muti-drug resistant HIV-1 infections. The studies described here elucidate the underlying mechanism by which the M66I substitution confers a marked resistance to the inhibitor. Furthermore, our structural findings will aid future efforts to develop the next generation of capsid inhibitors with enhanced barriers to resistance.

## INTRODUCTION

The human immunodeficiency virus 1 (HIV-1) core is a macromolecular complex formed during virion maturation. The outer shell of the native core is composed of viral capsid, which houses the viral RNA (vRNA) genome and key viral enzymes needed for infection. HIV-1 capsid contains ∼ 1,500 copies of capsid protein (CA), which in the presence of the cellular cofactor inositol hexaphosphate (IP) assemble into a closed conical structure composed of hexameric lattices and 12 pentamers (*1–3*). Key roles of viral capsid are to protect the vRNA genome from immune sensing during infection and enable the nuclear import of HIV-1 (*4–7*). The post-entry journey of the cores across the cytoplasm and into the nucleus is strongly influenced by host cell factors including the FG-motif-containing proteins CPSF6, NUP153, and SEC24C. These proteins bind to CA to enable trafficking of viral cores throughout the cytoplasm (SEC24C), import into the nucleus (NUP153), and subsequent intranuclear journey to preferred integration sites (CPSF6) (see (*8*) for a recent review).

Lenacapavir (LEN) is the first in class, highly potent and long-acting HIV-1 CA inhibitor, used to treat heavily treatment-experienced adults with multi-drug-resistant HIV-1 infection (*9–11*). LEN is administered through subcutaneous injections twice-yearly or taken once daily by mouth in combination with other HIV-1 medications. Furthermore, LEN is in Phase 3 PURPOSE 1 clinical trial for pre-exposure prophylaxis (PrEP) with interim results indicating 100% efficacy of subcutaneous LEN in protecting study participants from HIV-1 infection (*12*).

Cell culture assays demonstrated that LEN potently (EC_50_ of ∼50 - 100 pM) blocked replication of wildtype (WT) and multi-drug-resistant viruses that confer resistance to other retroviral drugs (*11, 13, 14*). The inhibitor impaired several capsid dependent HIV-1 replication steps. The highest potency of LEN was mapped to blocking the nuclear import and integration of HIV-1. The inhibitor hyper-stabilized HIV-1 cores in infected cells, thereby limiting the functionally essential pliability of mature CA lattices needed for the nuclear import and uncoating (*13, 15–17*). Higher LEN concentrations (∼ 10 - 100 nM) also impaired reverse transcription possibly through compromising the capsid integrity in the cytoplasm and exposing the vRNA genome to immune sensors (*15, 18, 19*). During virion maturation, LEN induced the accelerated, aberrant assembly of CA which resulted in inactive virus particles with eccentric morphology (*11, 14, 15*).

Structural and biochemical experiments demonstrated that LEN bound to CA hexamers with very high affinity and rigidified intra-and inter-hexamer interactions in the curved CA lattices (*11, 13*). The inhibitor primarily targets the crucial hydrophobic pocket of the CA N-terminal domain (*13, 20*), which also provides principal binding site for the FG-motif-containing cellular proteins CPSF6, NUP153, and SEC24C (*21–26*). LEN is a comparatively large molecule composed of four functional R1-R4 moieties (Fig. S1A). Particularly noteworthy is that the R3 difluorobenzyl ring, similarly to the aromatic Phe side chains from the cellular proteins, extends deep inside the pocket to form hydrophobic interactions with several CA residues including Met66. Yet, pharmacologically relevant LEN concentrations failed to displace CPSF6 from HIV-1 capsids (*27*). Tightly bound CPSF6 engaged with only a subset of hydrophobic CA pockets, thereby allowing LEN to access unoccupied hydrophobic pockets and hyper-stabilize the capsid (*27*).

Cell culture-based resistance selection experiments identified mutations in the vicinity of the LEN binding site including L56I, M66I, Q67H, K70N, N74D, Q67H/N74S, Q67H/T107N and Q67H/N74D substitutions (*11*). We previously elucidated the underlying mechanisms for resistance of Q67H, N74D, and Q67H/N74D variants to LEN (*28*). The Q67H substitution rendered a conformational switch, which adversely affected LEN binding; the N74D CA change resulted in the loss of hydrogen bonding and induced electrostatic repulsions between CA and LEN; and the Q67H/N74D CA variant exhibited cumulative effects of Q67H and N74D substitutions.

Emergent resistance associated mutations (RAMs) were also detected in clinical trials (*9, 10, 29*). In the Phase 2/3 CAPELLA study, 8 out of 72 patients, who received LEN to treat multi-drug-resistant HIV-1 infection, developed RAMs by week 26 (*10*). The detected substitutions in CA included M66I, M66I + N74D, M66I + Q67Q/H/K/N, M66I + T107A, Q67H + K70R, and K70H + T107T/N (*10*). Of these, the most prevalent was the M66I change present in 6 participants. Furthermore, of all RAMs detected in cell culture and clinical settings, the M66I substitution conferred by far the highest level of resistance to LEN (*11*).

Here, we report high resolution X-ray structures as well as biochemistry and virology studies of the M66I variant. Our findings revealed that the β-branched side chain of Ile66 introduced steric hindrance specifically to LEN. By contrast, the M66I substitution did not affect binding of FG-motif-containing cellular cofactors CPSF6, NUP153, or SEC24C to HIV-1 capsid. In cell culture, HIV-1_(M66I CA)_ failed to acquire compensatory mutations or replicate in the presence of LEN. Analysis of HIV-1 replication intermediates revealed that the mutant virus was severely impaired for nuclear import. Furthermore, the M66I substitution markedly altered HIV-1 integration targeting. Taken together, our findings provide mechanistic insights into how the M66I substitution confers remarkable resistance to LEN and influences HIV-1 replication. Moreover, our structural findings will aid future efforts to develop improved LEN analogs with enhanced resistance barriers.

## RESULTS

### The M66I mutation adversely affects binding of LEN but not FG-motif-containing cellular cofactors to CA hexamers

Met66 is an integral part of the hydrophobic pocket that provides the principal CA binding site for LEN and FG-motif-containing cellular proteins CPSF6, NUP153, and Sec24C. Therefore, we wanted to examine how the M66I change affected CA interactions with the inhibitor and the cellular cofactors. Binding of LEN to cross-linked CA hexamers was monitored by surface plasmon resonance (SPR), which allowed us to determine association rate (*k_on_*), dissociation rate (*k_off_*), and equilibrium dissociation (*K_D_*) constants (Fig. 1A, B). The M66I change did not significantly alter LEN *k_on_* values. By contrast, we observed about two orders of magnitude difference in *k_off_* values. While LEN remained tightly bound to WT CA hexamers during the course of the experiment (Fig 1A), the inhibitor rapidly dissociated from CA(M66I) hexamers (Fig 1B). Consequently, CA(M66I)_Hex_ exhibited a >230-fold reduced binding affinity for LEN compared to its WT counterpart.

**Fig. 1.**
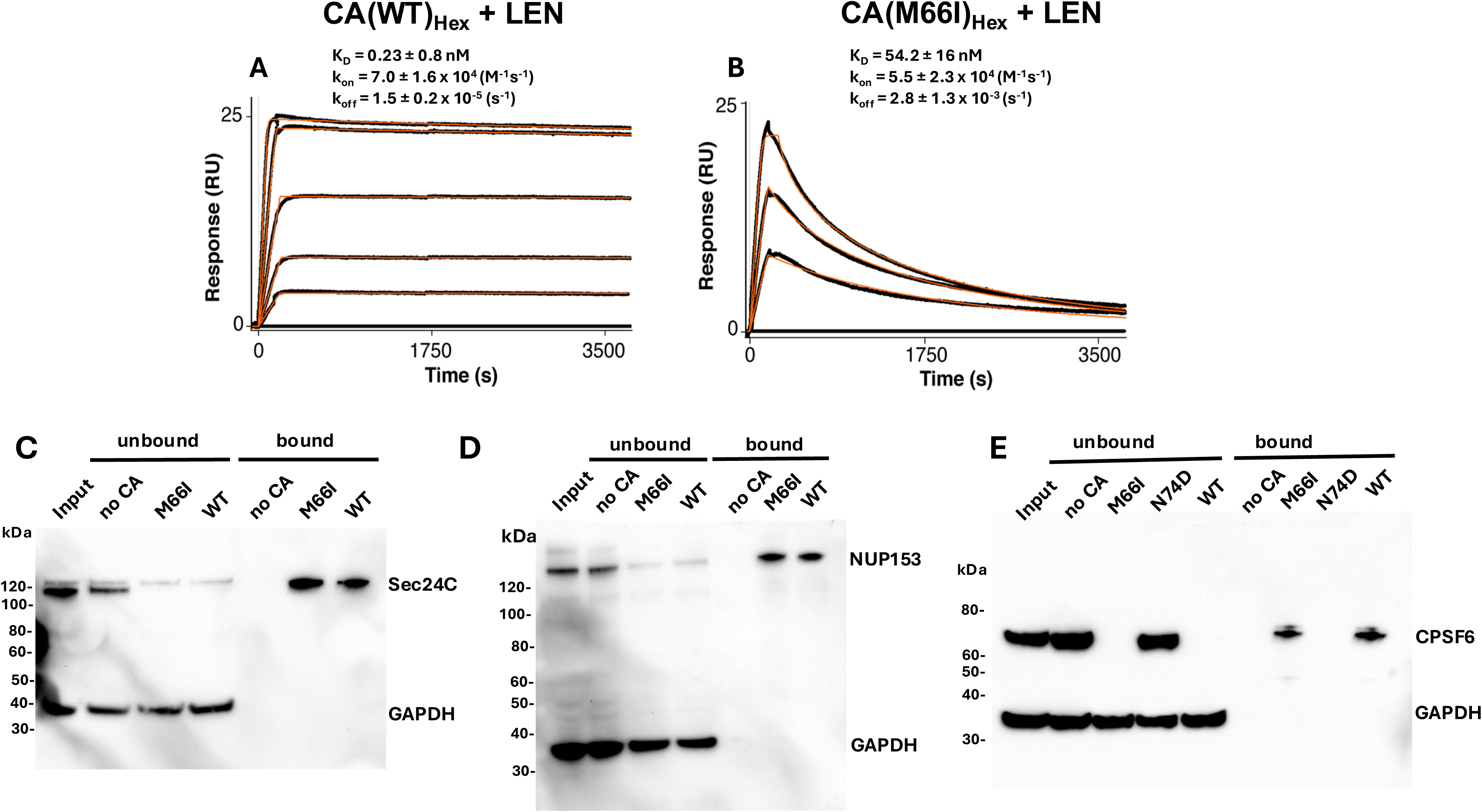
Effects of the M66I substitution on CA interactions with LEN and FG-motif-containing cellular cofactors. (A, B) Sensorgrams showing LEN binding curves to cross-linked WT (A) and M66I (B) CA hexamers. The response curves are shown in black, and the fit curves are shown in orange. The mean ± standard deviation (SD) equilibrium dissociation constant (*K_D_*), association rate constant (*k_on_*), and dissociation rate constant (*k_off_*) values were determined from three independent experiments with comparable results. (C-E) Co-pelleting experiments demonstrating binding of SEC24C (C), NUP153 (D), and CPSF6 (E) to WT and M66I nanotubes. GAPDH (panels C-E) and N74D nanotubes (panel E) served as CA-nonbinding negative controls.

To examine how the M66I change influences binding of FG-motif-containing cellular cofactors to CA, we tested the ability of nanotubes assembled from purified recombinant CA to co-pellet the cell factors from cellular lysates (Fig. 1C-E). In complete contrast to the inhibitor, the M66I change did not detectably affect binding of CPSF6, NUP153, or SEC24C to CA nanotubes. Effective binding of the full-length endogenous cellular proteins to both WT and M66I nanotubes was evident by depletion of these proteins from supernatants of cell lysates and their concomitant enrichments in the co-pelleted fractions (Fig. 1C-E). The N74D CA mutant, which is known to inhibit CPSF6 binding to CA (*30*), served as a control (Fig. 1E). Collectively, our biochemical studies have revealed that the M66I change specifically and markedly reduced LEN binding affinity to CA hexamers, whereas this mutation did not detectably affect capsid interactions with three different FG-motif-containing cellular cofactors.

### The M66I substitution creates steric hindrance specifically to LEN but not FG-motif-containing cellular cofactors

To understand the structural basis for the above biochemical findings (Fig. 1), we performed X-ray crystallography studies with cross-linked CA(M66I) hexamers. Our attempts to obtain a co-crystal structure of LEN bound to CA(M66I)_Hex_ were unsuccessful, likely due to the inability of the inhibitor to stably bind to the mutant protein (Fig. 1B). Instead, we obtained the structure of the unliganded CA(M66I) hexamer (Fig. 2, supplemental Table 1). To understand how the M66I change could impact LEN binding, we superimposed the structure of CA(M66I)_Hex_ onto WT CA_Hex_ + LEN. The results in Fig. 2A,B and Fig. S1 indicate that the β-branched side chain of Ile66 introduces steric hindrance with respect to the R3 and R4 rings of LEN. While van der Waals bonds usually exhibit carbon-carbon distances in the range of 3.3 -4.0 Å, the β-branched side chain (carbon γ2) of Ile66 was positioned within 2.5 Å and 2.7 Å of R3 and R4 groups, respectively. Accordingly, van der Waals surfaces shown in Fig. 2B indicate that the Ile66 side chain induced steric hindrance (see overlapping shaded areas) with respect to the aromatic rings of LEN. By comparison, the closest distances between Met66, which lacks the β-branched side chain, to R3 and R4 groups were 3.7 Å and 4.1 Å, respectively (Fig. 2C), indicating that Met66 in WT CA engages in hydrophobic interactions with LEN without encountering steric hindrance (Fig. 2D).

**Fig. 2.**
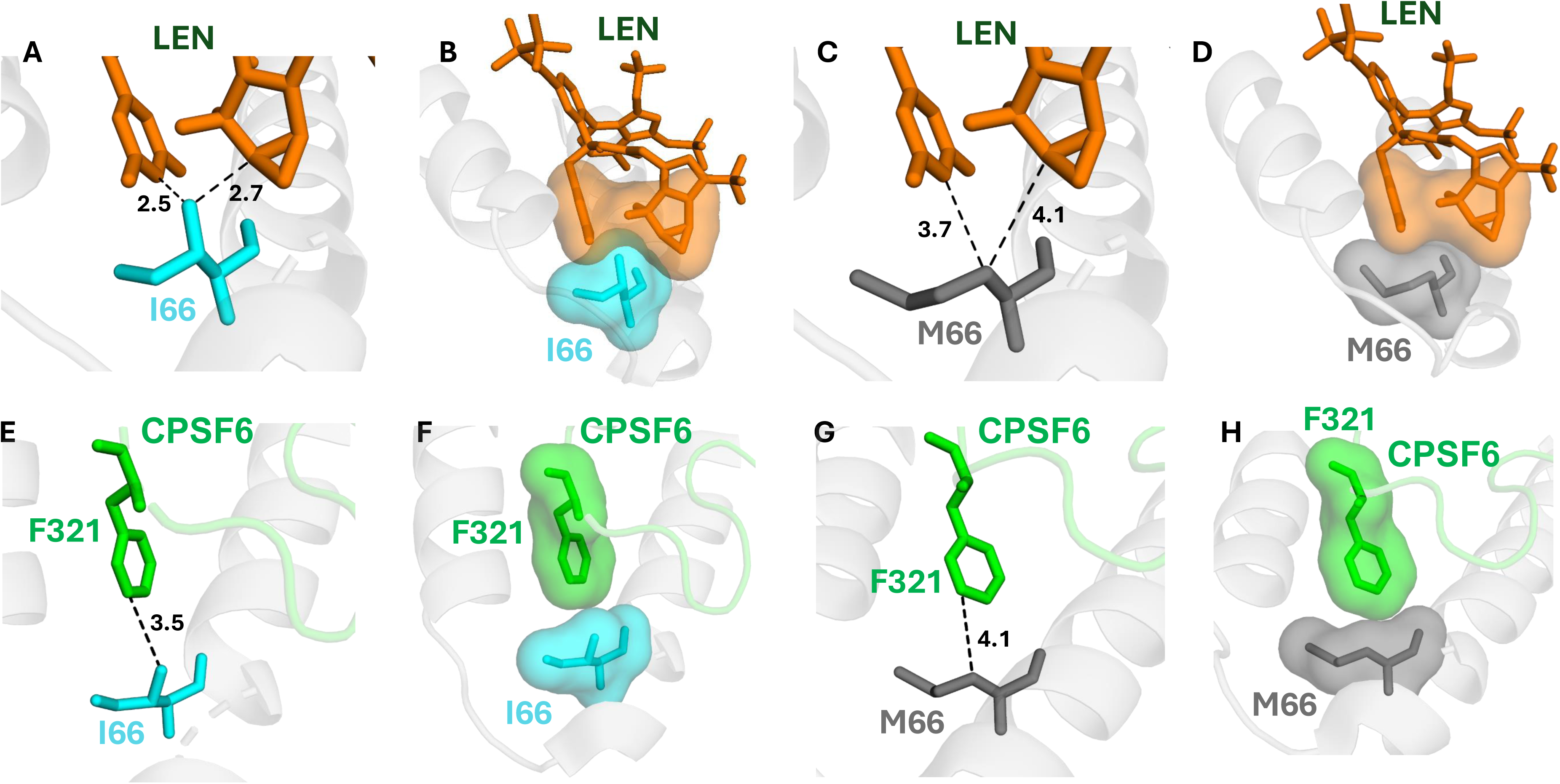
Structural bases for interactions of CA(M66I) hexamers with LEN and CPSF6. (A) The X-ray crystal structure of CA(M66I)_Hex_ illustrating the Ile66 side chain. The closest distances between the β-side chain carbon of Ile66 and R3 and R4 rings of LEN are indicated. (B) Van der Waals surfaces of the Ile66 side chain and LEN R3 and R4 groups are shown. The overlapping shaded areas indicate steric hindrance. (C) The X-ray crystal structure of WT CA_Hex_ illustrating the Met66 side chain. The closest distances between Met66 and R3 and R4 rings of LEN are indicated. (D) Van der Waals surfaces of the Met66 side chain and LEN R3 and R4 groups are shown to indicate mutual accommodation. (E) The co-crystal structure of CPSF6_313-327_ bound to CA(M66I)_Hex_. The closest distance between the β-side chain carbon of Ile66 and Phe321 side chain of CPSF6 is indicated. (F) Van der Waals surfaces of the CA Ile66 and CPSF6 Phe321 side chains are shown to indicate the lack of steric hindrance. (G) For comparison, the co-crystal structure of CPSF6_313-327_ bound to WT CA_Hex_ is shown. The closest distance between the Met66 side chain to the aromatic ring of the CPSF6 Phe321 side chain is shown. (H) Van der Waals surfaces of the WT CA Met66 and CPSF6 Phe321 side chains are shown to indicate lack of steric hindrance.

By contrast to LEN, we successfully co-crystalized the FG-motif-containing CPSF6_313-327_ peptide with CA(M66I)_Hex_ (Fig. 2E,F). The closest distance between the β-branched side chain (carbon γ2) of Ile66 to Phe321 of CPSF6 was 3.5 Å, which is within the range for hydrophobic interactions. Accordingly, steric clashes were avoided between van der Waals surfaces of CA Ile66 and CPSF6 Phe321 (Fig. 2F). For comparison we also show interactions of CPSF6 Phe321 to the WT CA hexamer (Fig. 2G,H). Collectively, our x-ray crystallography studies have revealed structural insights as to why the M66I change specifically affects binding of LEN but not CPSF6 to CA hexamers.

### HIV-1_(M66I CA)_ fails to replicate or acquire compensatory mutations in the presence of LEN during propagation in cell culture

To assess how the M66I change influences viral replication kinetics and select for potential compensatory mutations, we transfected SupT1 cells with a full-length NL4-3 infectious molecular clone (Fig. S2). While replication of the WT virus peaked at day 10 post-transfection, HIV-1_(M66I CA)_ was generally unable to replicate over several weeks of propagation in culture. Occasionally, we observed delayed replication of the mutant virus (Fig. S2). Sequencing of genomic DNA isolated from infected cells revealed that replication resulted from the acquisition of a mutation, reverting Ile66 to Met66. The reversion of the mutant virus to its WT counterpart likely occurs because the Met-to-Ile amino acid substitution is conferred by a single nucleotide change, providing M66I with a simple pathway back to full replicative capacity. To test whether LEN can prevent M66I reversion to WT during propagation, we performed similar experiments in the presence of the inhibitor (Fig. 3). For this, we infected SupT1 cells with WT or M66I virus in the presence of DMSO (Fig. 3A), 1 nM LEN (Fig. 3B), or 10 nM LEN (Fig. 3C). Similar to the transfection experiment, we occasionally observed M66I reversion to WT in the absence of LEN (Fig. 3A). However, HIV-1_(M66I CA)_ failed to replicate in the presence of LEN (Fig. 3B, C), suggesting that HIV-1 is unable to adapt to the M66I-induced fitness defect via acquisition of second-site compensatory mutations. The lack of M66I variant reversion to WT in the LEN-containing cultures was likely due to the ability of the inhibitor to fully suppress reverted WT viral replication.

**Fig. 3.**
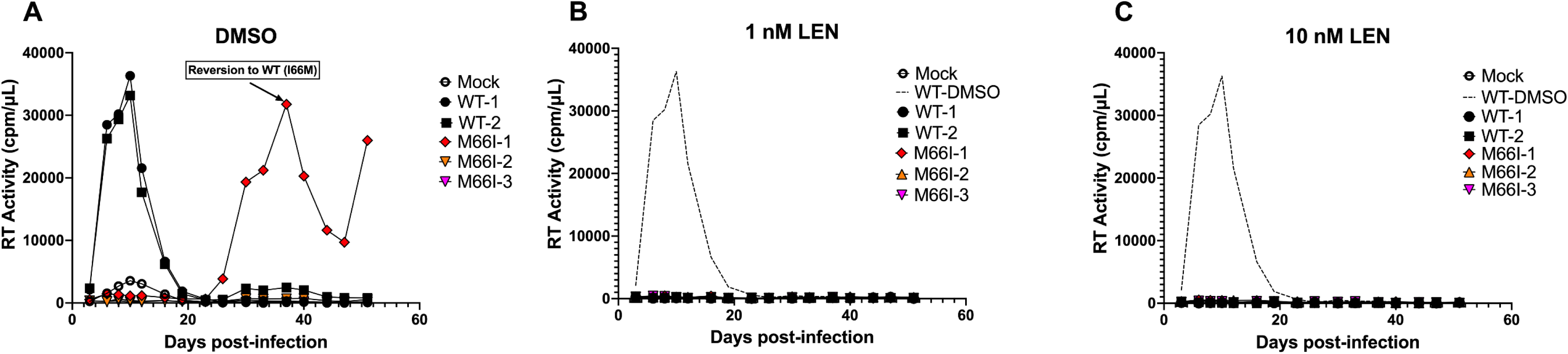
HIV-1_(M66I CA)_ does not replicate in the presence of LEN. SupT1 cells were infected with NL4-3 WT or M66I virus in the presence of DMSO (A), 1 nM LEN (B), or 10 nM LEN (C). Replication kinetics were assessed by quantifying supernatant RT activity at 2 - 4-day intervals for ∼50 days post-infection. Data are representative of three independent experiments.

### Maturation of HIV-1 particles is marginally affected by the M66I change

Our follow up mechanistic studies have focused on elucidating the distinct steps of HIV-1 replication cycle that could be adversely affected by the M66I mutation. For this, we initially examined late steps of HIV-1 replication. Fig. S3A shows that the M66I mutation did not substantially affect Gag expression in HEK293T cells transfected with full-length WT or M66I molecular clones. Extracellular levels of virions were also comparable for WT and M66I viruses (Fig. S3B). Immunoblotting analysis of isolated virions indicated that proteolytic processing of Gag and Gag-Pol polyproteins were not affected by the M66I change, as evidenced by similar levels of CA p24, matrix p17, and reverse transcriptase (RT) p66/p51 observed in WT and mutant virions (Fig. S3C).

Next, we analyzed morphologies of HIV-1 particles using transmission electron microscopy (TEM) (Fig 4A-B). Distinct species were observed for both WT and M66I virus particles including mature, immature, and atypical structures. The atypical species included empty cores, rod-shaped cores, multiple cores, multiple densities, and empty cores, which were grouped as cumulative totals of defective or atypical virions. The mature particles were predominant in both WT (∼82%) and M66I (∼64%) samples. A relatively modest reduction of mature virions conversely correlated with a concomitant increase in atypical species for HIV-1_(M66I CA)_ compared to its WT counterpart. Both WT and mutant virus samples contained only residual amounts of immature virions, thereby providing an additional line of evidence that the M66I change did not affect proteolytic processing of Gag.

**Fig. 4.**
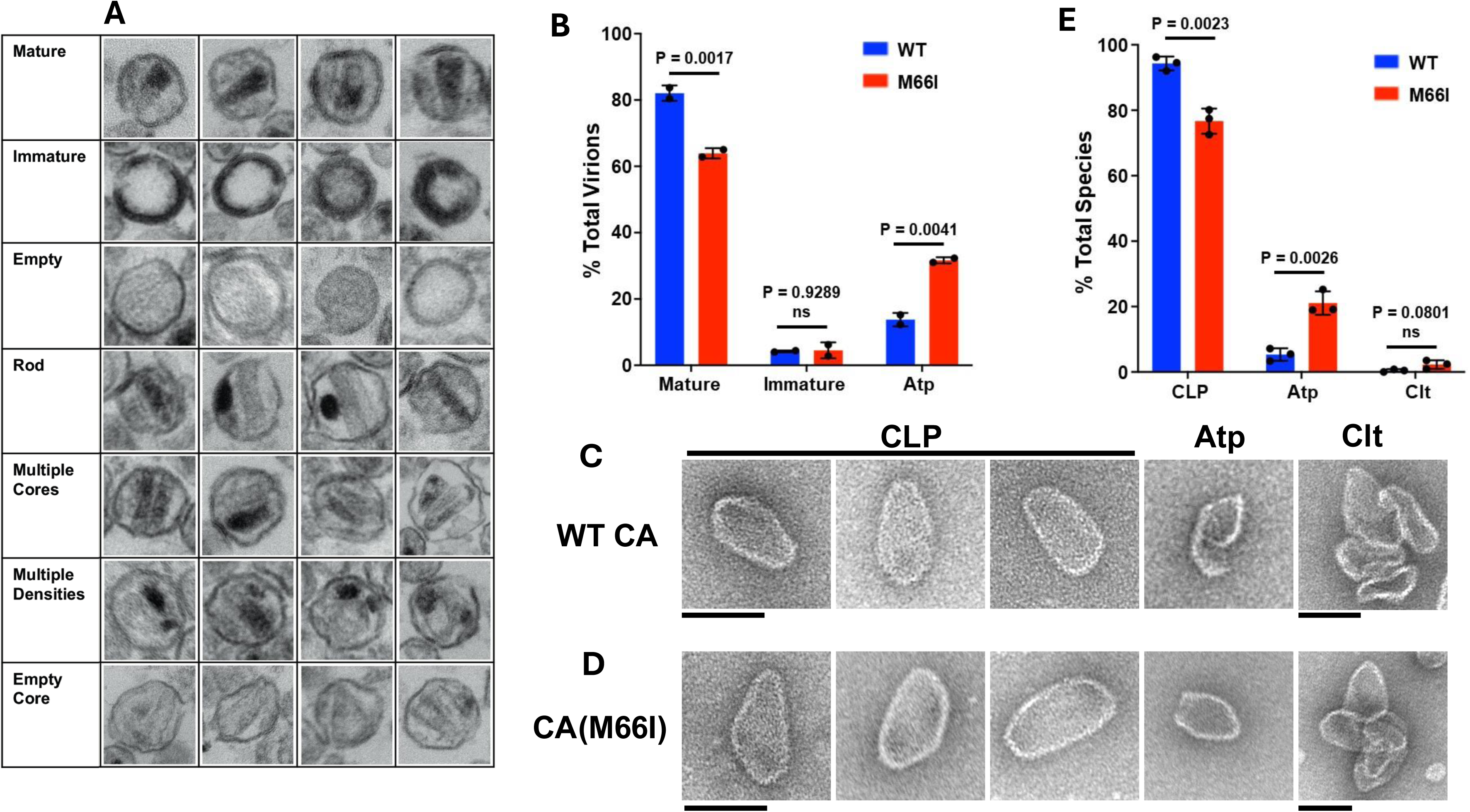
TEM analysis of HIV-1_(M66I CA)_ native cores and CLPs assembled with CA(M66I). (A) Representative TEM images of virions used during morphological classification. For each experimental condition, >200 virions were counted. Across cumulative experiments (n=2 for WT and M66I), >520 virions were counted in total. Virions with visibly electron-dense outer membranes were classified as: mature, containing a single circular or conical electron-dense region; immature, containing outer, comparatively thick toroidal or semi-hemispheric electron density; empty, comparative lack of luminal contents; rod, electron-dense or electron-lucent core lacking obvious conical shape; multiple cores, >1 conical, circular, or rod-like capsid shape that was electron-dense or lucent; multiple densities; >1 distinct electron-dense region wherein at least one of the densities could not be readily attributed to a core structure; empty core, roughly conical shapes that lacked associated electron density. (B) Percentage of virions analyzed by TEM belonging to various morphological categories (avg. ± SEM for n=2 independent experiments). P values for statistical differences between samples are shown. ns, not significant (P>0.05). (C, D) In vitro, IP6 mediated assembly of CLPs from WT (C) and M66I (D) CAs. Representative micrographs are shown. (E) Quantification of the CLP assembly products. The averaged data (+/− SD) from three independent experiments are shown. CLP, capsid-like particles. Atp, atypical assemblies. Clt, clusters. Scale bars: 100 nm.

For complementary biochemical experiments, we visualized capsid like particles (CLPs) assembled from purified recombinant WT and M66I CAs in the presence of IP6 (Fig. 4C-E). Akin to WT CA, CA(M66I) predominantly yielded conically shaped CLPs. In fact, these in vitro experiments (Fig. 4C-E) exhibited a similar trend as observed with native virions (Fig. 4A-B); CLP levels were only modestly decreased with concomitant increases of the atypical structures for CA(M66I) vs WT CA. Taken together, our virology and biochemistry experiments indicated that M66I change only modestly affected formation of the mature capsid.

### The M66I substitution enhances the stability of HIV-1 cores

We next examined how the M66I change influenced the stability of native viral cores using the previously described CypA-DsRed (CDR) loss assay (*13, 31*). Towards this goal, we co-packaged WT and M66I mutant viruses with the Vpr-Integrase-mNeonGreen (INmNG) and CDR marker proteins. Analysis of *in vitro* capsid/CDR-marker loss following virus membrane permeabilization with a mild detergent (Saponin, 100 μg/ml) revealed a substantial increase in the stability of M66I cores compared to their WT counterparts (Fig. 5A). The kinetics of 50% CDR loss from the M66I variant was substantially delayed (*t_1/2_* ∼12.5 min) compared to WT cores (*t_1/2_* ∼6.5 min), resulting in a >2-fold increase in the number of stable cores detected at the end of 25 min imaging (Fig. 5A). Similarly, analysis of capsids that productively entered the cytoplasm of living cells showed a ∼3-fold increase in the stability of M66I cores compared to the WT cores. Taken together, our results suggest that M66I substitution enhances the stability of HIV-1 cores.

**Fig. 5.**
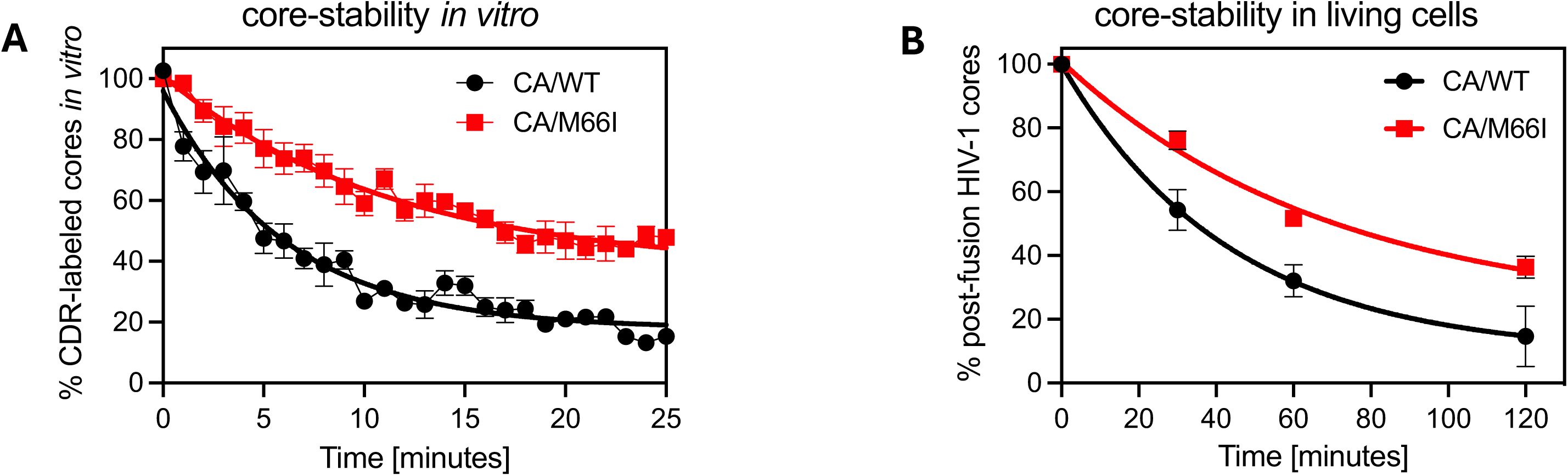
HIV-1_(M66I CA)_ cores are more stable than their WT counterparts. Quantification of the avid CDR/capsid marker loss from INmNG labeled cores *in vitro* (A) and inside TZM-bl cells (B) demonstrating that M66I substitution enhances the stability of viral cores. Error bars are mean and SEM from 4 independent experiments.

### HIV-1_(M66I CA)_ is impaired for the penetration of the nuclear pore complex (NPC) and nuclear import

To investigate how the M66I change affects early steps of HIV-1 infection, we examined replication intermediates in target cells. Relative vRNA levels for the WT virus and HIV-1_(M66I CA)_, assessed at 2 h post-infection, were comparable (Fig. 6A), indicating that the mutation did not influence viral entry into target cells. While the RT enzyme in the mutant virus (Fig. 6B) was fully active, synthesis of reverse transcription products at 12 h post-infection was reduced by ∼3-fold (Fig. 6C). Levels of 2-LTR circles, which are a surrogate marker for nuclear import, were markedly decreased for the mutant virus (Fig. 6D), suggesting a severe nuclear import defect.

**Fig. 6.**
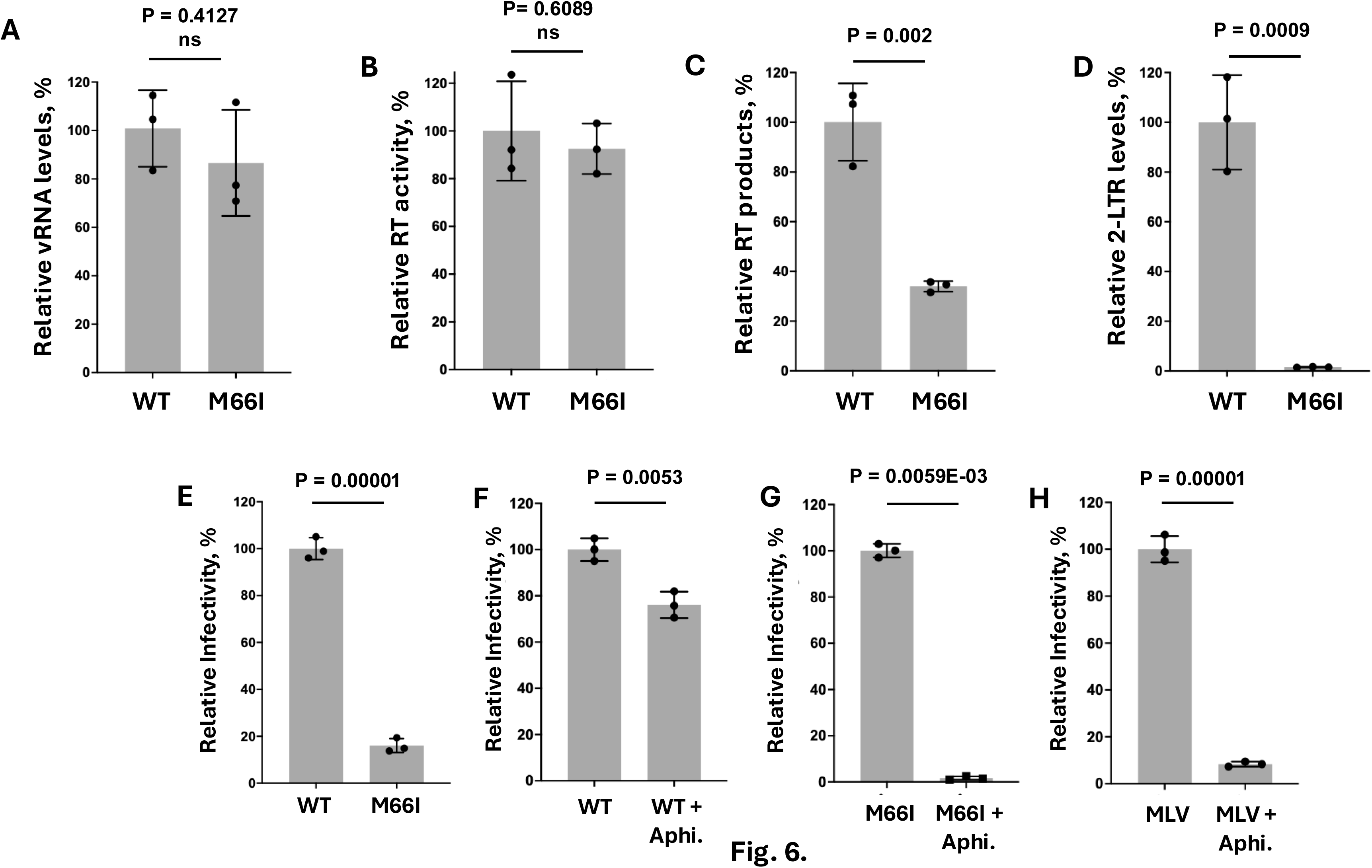
HIV-1_(M66I CA)_ is severely impaired for nuclear import. Analysis of HIV-1 replication intermediates during the early phase of virus replication. (A) Viral RNA levels post-entry. MT4 cells (2 × 10^5^) were infected with VSV-G pseudotyped HIV-1 (500 ng of p24) followed by incubation, first on ice for 15 min and then at 37 °C for 2 h. Cells were washed four times with PBS and then treated with 0.25% trypsin to remove unbound viruses. Total RNA was isolated and quantified via RT-qPCR. (B) Reverse transcriptase activity. VSV-G pseudotyped viruses were produced by transfection and quantified by p24 ELISA. RT activity was measured using a colorimetric assay with 10 ng of p24 HIV particles. (C-E) Reverse transcription products (C), 2-LTR circles (D), and infectivity (E) during HIV-1 single-round infection. 500 ng of p24 particles were used for infection in (C, D), while 10 ng of p24 particles were used for infectivity in (E). (F-H) Effects of aphidicolin on single-round infection of WT HIV-1, HIV-1_(M66I CA)_, and MLV. Cells were treated with 1 µg/ml aphidicolin or DMSO for 24 h, then reseeded with or without aphidicolin (1 µg/ml) and infected with 10 ng of p24 WT particles (F), 100 ng of HIV-1_(M66I CA)_ particles (G), or 100 µl of MLV particles. Infectivity was assessed 48 hpi by measuring the luciferase activity. Data (mean ± s.d.) from three independent experiments are shown. P values for statistical differences between samples are shown. ns, not significant (P>0.05).

To further characterize the effect of the M66I mutation on nuclear import, we compared the infectivity of HIV-1_(M66I CA)_ with the WT virus in the absence and presence of the cell division inhibitor aphidicolin (Fig 6E-G). In a single-round infectivity assay, HIV-1_(M66I CA)_ was ∼10-fold less infectious than its WT counterpart (Fig. 6E). As expected (*32*), in control experiments, aphidicolin treatment only modestly decreased WT HIV-1 infectivity (Fig. 6F), which is consistent with the ability of WT cores to traverse through the NPC into the nucleus of non-dividing cells. By contrast, aphidicolin treatment markedly impaired the infectivity of HIV-1_(M66I CA)_ (Fig. 6G) and the control murine leukemia virus (MLV) (Fig. 6H), which is dependent on cell division for its entry into the nucleus (*33*). These results further highlight that the block to HIV-1_(M66I CA)_ infectivity is due to its inability to traverse through the NPC of aphidicolin-arrested cells.

To gain additional insights into the observed nuclear entry block of the M66I variant, we employed live-cell confocal imaging to visualize INmNG-labeled viruses in aphidicolin-arrested TZM-bl cells. An extended imaging (∼16 h) and unbiased single virus tracking approach (*17*) were employed to spatiotemporally resolve and quantify multiple virus entry steps, including (1) docking interactions between HIV-1 and the nuclear envelope (NE), (2) nuclear entry of the INmNG-cores, and (3) infection, measured by the expression of an eGFP-reporter from integrated viral DNA.

Single virus trajectory analysis revealed that similar to the WT virus, HIV-1_(M66I CA)_ cores retained the ability to reach and stably dock at the NE (Fig. 7A-C). A similar proportion (∼40%) of both WT and M66I INmNG labeled cores docked at the NE (Fig. 7B) as defined by the localization of a single INmNG-core within a 360 nm radius (2 pixels) on the NE for longer than 7.5 min (*17*). The docking events for both WT and M66I cores were significantly enriched compared to sham infections conducted with virions assembled in the absence of vesicular stomatitis virus glycoprotein G (no-VSV-G; 15.8%), which defined non-specific interactions between the NE and endosomal resident INmNG-cores that entered the cell without endosome escape into the cytosol (Fig. 7B). Cumulative analysis additionally showed that ∼30% of M66I and ∼26% of WT cores remained docked on the NE for longer than 10 min. This long-docking profile was significantly higher (>5-fold) than a few non-specific docking events detected with the no-VSV-G control (5.1%, Fig. 7C). Compared to WT cores, which retained the ability to enter the nucleus for infection (Fig. 7D), M66I cores notably docked with the NE for long-durations (lasting up to 60 min, Fig. 7C), but failed to enter the nucleus. These results are fully consistent with the notion that M66I cores are severely compromised for the passage through the NPC.

**Fig. 7.**
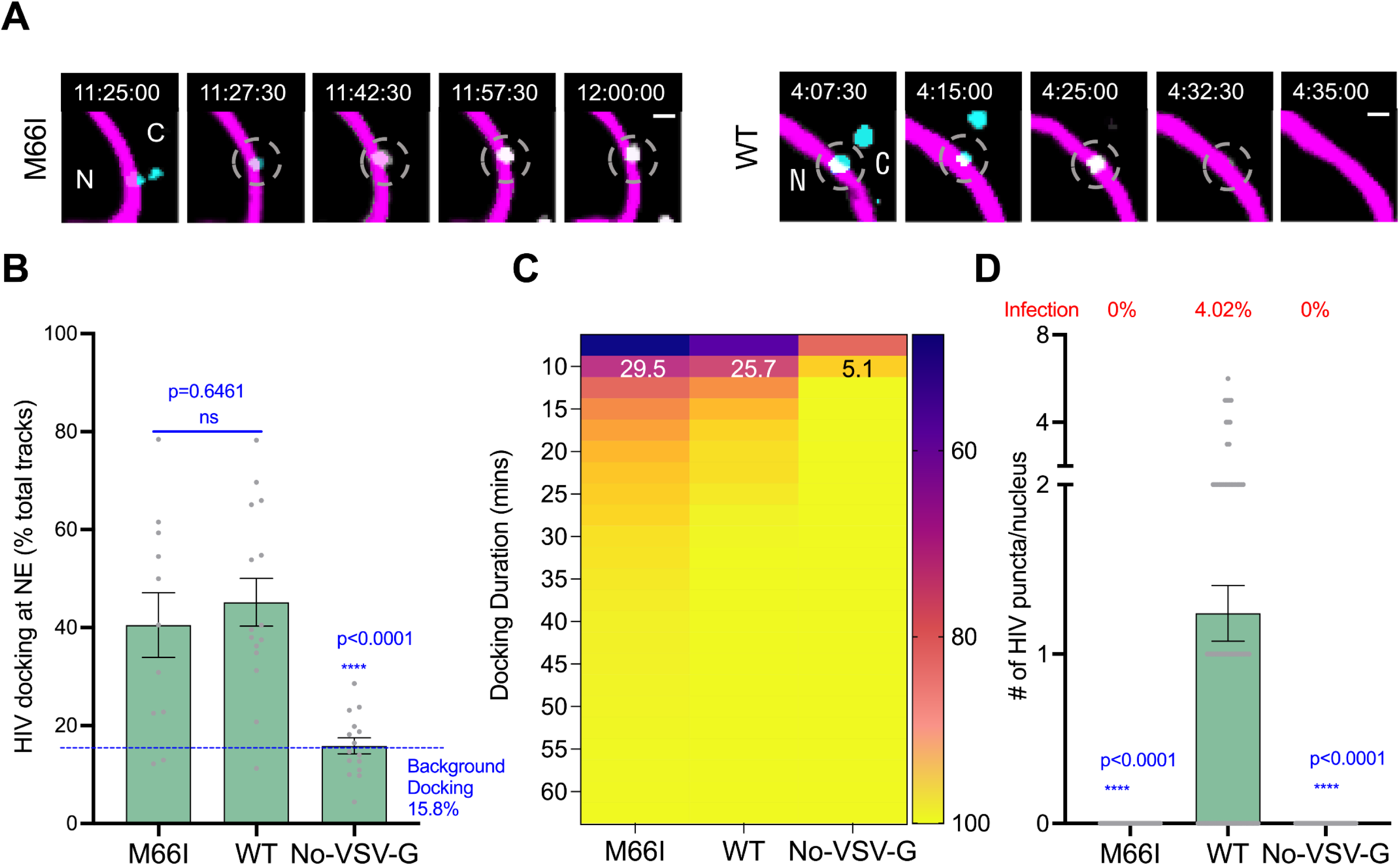
HIV-1_(M66I CA)_ cores reach and dock at the nuclear membrane, but fail to enter the nucleus. (A) Representative single z-stack time-series images showing interactions between M66I (*left panels*) and WT (*right panels*) cores with the NE. The INmNG labeled HIV-1 cores (*cyan, puncta*) and the nuclear lamina (*magenta*) are shown. The segments of HIV-1 docking at the NE are marked by dashed circles. Time stamps, and locations of cytoplasm (C) and nucleus (N), are superimposed in images. The scale bar is 1 µm. (B) Quantification of docking events in individual cells (*grey dots*), and (C) the cumulative frequency plot of the duration of docking demonstrating that both WT and M66I cores reach and dock at the NE with the similar frequency, which are significantly higher than the non-specific No-VSV endosomal interactions. Docking events lasting for longer than 10 min are superimposed on top of the heat map in (C). (D) Quantification of the number of HIV-1 INmNG puncta that entered the nucleus during the first 8 hpi demonstrating that only WT, but not M66I, cores enter the nucleus. The % of eGFP-reporter expressing infected cells at the end of 16 h of the movie is overlaid in (D). Error bars are mean, and standard errors are from 10 nuclei in (B) and the entire nucleus volume of >100 nuclei in (D). Student’s ‘t’-test was used to determine statistical significance in (B and D).

### The M66I substitution retargets HIV-1 integration

Mutations that compromise nuclear import are known to alter HIV-1 integration site distribution (ISD) (*34*). Accordingly, we asked if the M66I change affected the ISD of HIV-1. The WT virus favors integrating into speckle associated domains (SPADs) and disfavors lamina associated domains (LADs) (*35–37*). This pattern was markedly altered by the M66I variant. HIV-1_(M66I CA)_ integration into SPADs (∼6%) was substantially reduced compared to the WT levels (∼31%) and trended closer to the random integration control (RIC, Fig. 8A). Conversely, the M66I variant integrated into LADs more frequently than WT virus (Fig. 8B). The WT virus is also known to preferentially integrate into active transcription units and gene dense regions. Integration into genes was reduced from ∼83% to ∼70% for the CA mutant virus (Fig. 8C), and from ∼21 genes/Mb to ∼11 genes/Mb (Fig. 8D). Furthermore, the M66I variant integrated less frequently into CpG islands and transcription-start sites than its WT counterpart (Fig. 8E,F). Collectively, these results demonstrate that the M66I change resulted in marked retargeting of HIV-1 integration.

**Fig. 8.**
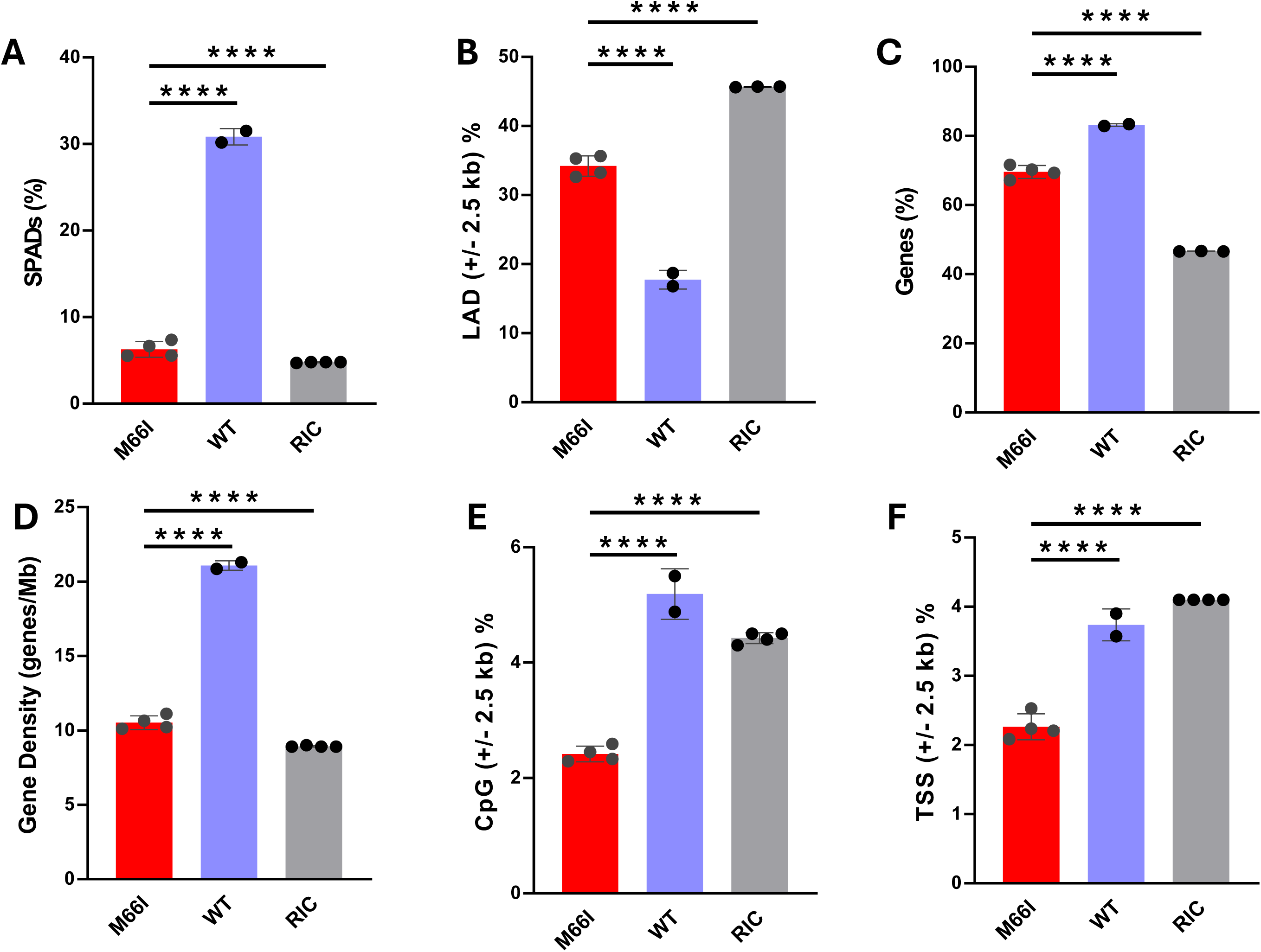
The M66I substitution alters HIV-1 integration targeting. HEK293T cells were infected with VSV-G pseudotyped WT HIV-1 and HIV-1_(M66I CA)_. 5 days later cells were collected, and genomic DNA was extracted and used for LM-PCR and Illumina sequencing. Distribution of HIV-1 integration sites for HIV-1_(M66I CA)_ and the WT virus in SPADs (A), proximal to LADs (B), in genes (C), gene dense regions (D) and proximal to CpG islands (E) and transcriptional start sites (TSSs) (F) are shown. Random integration (RIC) is included for comparison. P values (****p<0.00001) were calculated by Fisher’s exact test or by Wilcoxon sum rank test (for Gene density) and are reported in supplementary table 2.

## DISCUSSION

Our structural studies uncovered the underlying mechanism for how the M66I variant confers a remarkable resistance to LEN. Unlike Met66, which establishes crucial hydrophobic interactions with the difluorobenzyl (R3) and the cyclopenta-pyrazole (R4) groups of LEN, the β-branched side chain of Ile66 induced steric hindrance with respect to both R3 and R4 rings. Consequently, the M66I change dramatically increased the dissociating rate (*k_off_*) of the inhibitor from CA hexamers and correspondingly decreased the binding affinity.

Unexpectedly, the FG-motif-containing cellular cofactors that bind to the same hydrophobic pocket of CA as LEN were unaffected by the M66I change. CPSF6 Phe321, similarly to the R3 group of LEN, extended deep inside the hydrophobic CA pocket. Nevertheless, unlike LEN, our structure of CPSF6_313-327_ bound to CA(M66I)_Hex_ showed the lack of steric hindrance between CA Ile66 and CPSF6 Phe321. Instead, the β-branched side chain of Ile66 established hydrophobic interactions with the aromatic ring of Phe321. Consistent with these structural observations, our biochemical experiments demonstrated that the M66I change did not affect binding of cellular CPSF6 to CA nanotubes. Similarly, other FG-motif-containing cellular cofactors NUP153 and SEC24C bound with similar efficiencies to WT and M66I nanotubes. Taken together, our findings indicate that the M66I change specifically and adversely affected binding of LEN to HIV-1 capsid, whereas this substitution did not influence capsid interactions with cognate FG-motif-containing cellular cofactors.

A recent structural study identified Met66 as a crucial gate keeper amino acid which regulates the formation of essential capsid assembly intermediates hexamers and pentamers (*20*). Specifically, the Met66 side chain, which extends from the H4 helix, influences the secondary structure of the C-terminal end of the H3 helix (*20*). In the hexamers, this segment adopts disordered conformation to accommodate the extended side chain of Met66 (Fig. S4A), whereas in the pentamers the Met66 side chain is repositioned to allow the elongated α-helical structure of H3 (Fig. S4B). It was shown that the M66A change markedly altered the assembly process by yielding small spheres composed exclusively of CA pentamers (*20*). These observations could be explained by the inability of a short side chain of Ala66 to reach the H3 to alter its α-helical conformation (Fig. S4C). Unlike Ala66, the longer and bulkier side chain of Ile66, similarly to Met66, extends further toward the H3 (Fig. S4D) and could regulate the conformational switch needed for formation of both hexamers and pentamers. Indeed, in sharp contrast from M66A, the M66I variant predominantly yielded mature, conically shaped capsids, albeit at marginally reduced levels compared to the WT counterparts (Fig. 4 and Fig. S4E-G).

Analysis of HIV-1 replication intermediates indicated that the most profound effect of the M66I substitution was to impair HIV-1 nuclear import. Consistent with our biochemical data demonstrating the ability of CA(M66I) to retain WT levels of binding to the FG-motif-containing cellular cofactors, HIV-1_(M66I CA)_ cores stably bound to the NE in infected cells. However, the mutant cores failed to penetrate inside the nucleus. The underlying mechanism(s) for these observations is unclear. Previous studies demonstrated the importance of capsid pliability for HIV-1 nuclear import (*13, 38, 39*); the substitutions that either stabilized or destabilized the capsid lattice impeded the cores to enter the nucleus. Our findings indicate that the M66I capsid exhibited higher stability than its WT counterpart, which in turn could adversely affect the M66I variant to penetrate through the NPC. We furthermore speculate that the enhanced rigidity coupled with the inability to fully enter inside the nucleus could trigger premature uncoating of HIV-1_(M66I CA)_ capsid at the NPC, which would be consistent with observed retargeting of integration sites into LADs located adjacent to the NE. In turn, enhanced integration frequency in the transcriptionally repressed LAD sites warrants further investigation into a potential role of the clinically relevant M66I variant in establishing latent viral reservoirs.

Our cell culture experiments demonstrated that HIV-1_(M66I CA)_ failed to acquire compensatory mutations. In the absence of LEN, the mutant virus reverted to its WT counterpart, whereas in the presence of pharmacologically relevant concentrations of the inhibitor sufficient to fully suppress reverted WT viruses, HIV-1_(M66I CA)_ failed to replicate. These findings have potential clinical implications, as they indicate that the ability of the M66I variant to confer marked resistance to LEN is countered by the inability of HIV-1_(M66I CA)_ to acquire compensatory mutations or effectively replicate in the presence of the inhibitor. We also note that while the M66I substitution was a prevalent RAM that emerged in the background of multi-drug-resistant HIV-1 phenotypes in the Phase 2/3 CAPELLA study (*9, 10*), in a separate phase 2 CALIBRATE trial, where LEN was used in combination with other antiretroviral therapies to treat adults who had not previously received antiretrovirals, emergent RAMs were found in only two of 249 participants: one containing Q67H and another Q67H/K70R substitutions in CA (*40*). These limited data do not allow us to conclude whether the M66I substitution is more likely to develop in the background of multi-drug-resistant viruses or whether it can also emerge with prolonged clinical use of LEN in treatment naïve individuals. Future mechanistic studies are warranted to characterize the M66I variant in the context of multi-drug-resistant viral phenotypes.

Taken together, this study provides mechanistic insights into how the M66I substitution confers remarkable resistance to LEN and affects HIV-1 replication. Furthermore, our structural findings will aid future medicinal chemistry efforts to rationally develop improved LEN analogs with enhanced barriers to the generation of resistance.

## MATERIALS AND METHODS

### Preparation of purified recombinant proteins

WT CA and CA_(A14C/E45C/W184A/M185A)_ were used for preparation of monomeric proteins and cross-linked hexamers, respectively. The M66I mutation was introduced in pET3a using a QuikChange XL site-directed mutagenesis kit (Agilent). Recombinant proteins were expressed in *E. coli* (BL21-DE3) cells and purified as previously described (*41–44*). Briefly, (NH_4_)_2_SO_4_ was used for initial precipitation of the protein followed by two column chromatography steps with HiTrap Q-Sepharose High Performance 5 mL column (Cytiva) and Cytiva HiLoad 16/600 Superdex 200 pg column. The disulfide-stabilized WT and M66I hexamers were assembled, purified and concentrated as reported (*28*).

### SPR

SPR experiments were performed using a 4-channel Reichert 4-SPR equipment. A nitrilotriacetic acid (NTA) sensor chip was conditioned with 40 mM NiSO_4_ at a flow rate of 50 μL/min for 3 min. Cross-linked His_6_-CA hexamers (WT and M66I) were immobilized on the NTA sensor chip via their C-terminal His-tags. The running buffer contained 0.01 M HEPES (pH 7.4), 0.15 M NaCl, 0.05% vol/vol Surfactant P20, and 5% DMSO. LEN was prepared as described (*28*). The sensor chip was regenerated with 350 mM EDTA and 50 mM NaOH. For each interaction, one channel was used as a control to account for background binding and drifting. Data were analyzed using Scrubber 2.0 and TraceDrawer and fit with a simple kinetic model with a term for mass transport. The mean ± standard deviation (SD), equilibrium dissociation constant (*K_D_*), association rate constant (*k_on_*), and dissociation rate constant (*k_off_*) values were determined from three independent experiments with comparable results.

### Co-pelleting assay

WT and M66I nanotubes were assembled by incubating 42.5 µM protein in a buffer containing 25 mM Tris-HCl, pH 8.0 and 2 M NaCl overnight on ice. Cellular lysates were added to the preformed CA nanotubes and incubated for 30 min at 4 °C. The mixtures were centrifuged at 21,000g for 15 min at 4 °C. Supernatants and pellets were analyzed by SDS–PAGE and immunoblotting.

### Assembly and TEM analysis of CLPs

Conical HIV-1 CLPs were assembled by incubating purified recombinant CA (3 mg/mL) with 500 μM IP6 (Sigma), 50 mM MES, pH 6, 40 mM NaCl, and DMSO at room temperature for 16 h. CA protein assembly reactions were applied to glow-discharged formvar/carbon 300-mesh Cu grids for negative staining and TEM imaging. Assembly samples (10 μL) were applied to glow-discharged formvar/carbon 300-mesh Cu grids (Electron Microscopy Sciences, Hatfield, PA) and incubated for 2 min. After blotting, grids were washed for 2 min with 0.1 M KCl. Subsequent blotted grids were stained with 2% uranyl acetate for 2 min and blotted dry. Micrographs were collected by FEI Tecnai T12 Spirit (ThermoFisher, Waltham, MA) at 120 kV.

### X-ray crystallography

Apo crystals of CA(M66I)_Hex_, and co-crystals of CA(M66I)_Hex_+IP6+CPSF6_313-327_, were grown via hanging drop vapor diffusion at 4°C with an equal volume of crystallization buffer. The crystallization buffers for CA(M66I)_Hex_ crystals contained 0.125-0.35M sodium iodide, 2-8% Peg 3350, 6% glycerol, and 0.1 M sodium cacodylate pH 6.5. For the CA(M66I)_Hex_+IP6+CPSF6_313-327_ crystals, the buffer contained 0.1 M Tris hydrochloride pH 7.8, 11% PEG 8000 and 5% glycerol, with 5 mM of IP6 and 2 mM of CPSF6_313-327_ was added to the protein droplet. The cryogenic solutions for the crystals consisted of crystallization buffer with additional 20% glycerol. Crystals were flash frozen in liquid nitrogen and the data were collected at the Advanced Light Source, Beamline 4.2.2 (Macromolecular Crystallography; MBC) at 100 K and at a wavelength of 1.00003 Å. Data reduction and scaling was performed using XDS (*45*). PHASER (*46*) from the PHENIX suite (*47*) was used to perform molecular replacement, using PDB 3H47 as a search model for CA(M66I)_Hex_ and 7SNQ as a search model for the tripartite complex. The structures were refined using repeated cycles of model building and refinement via COOT (*48*) and phenix.refine (*47*), respectively. Ligands were individually oriented into the structures based on the Fo-Fc omit map density at 3 σ and were then refined with phenix.refine to ensure that they fit the 2Fo-Fc density at 1 σ (*48*). Water molecules were originally added through phenix.refine and then evaluated individually to ensure that they fit the 2Fo-Fc density at 1σ (*49*). The quality of the final model was assessed using Molprobity (*50*). The coordinates are deposited in the Protein Data Bank under accession codes 7RAO and 8GDV. The data collection and refinement statistics are given in Table S1.

### Virion production and analysis of viral replication intermediates

Luciferase-encoding pseudotyped HIV-1 viruses were produced by co-transfecting HEK293T cells with VSV-G and pNL4-3-E-R-Luc plasmids. Lentiviral vectors were produced by co-transfecting VSV-G, Δ8.2, and Tsin-IRESpuro based plasmids in HEK293T cells. The MLV gammaretroviral vector was produced by co-transfecting VSV-G, PJK3, pL-VSVG and pCMV-tat in PhoenixAMPHO cells (*24*).

HIV-1 RNA levels were analyzed by RT–qPCR as previously described (*24, 27*). Virions produced from HEK293T cells were quantified by HIV-1 P24 ELISA (ZeptoMetrix) and treated with 60 U/ml of DNase I (Ambion) to remove the residual plasmid. MT4 cells (2 × 10^5^) were infected with VSV-G pseudotyped HIV-1 virions (500 ng of p24) followed by incubation, first on ice for 15 min and then at 37 °C for 2 h. The cells were washed with PBS four times and then treated with 0.25% trypsin to remove the unbound virus. Trypsin was inactivated by adding culture medium containing 10% FBS and the cells were subsequently washed three times. RNA was isolated by using RNeasy Mini Kit (Qiagen). Quantification of vRNA was performed by using reverse transcription and real-time PCR of conserved sequences within Gag as described (*51*).

Quantification of HIV reverse transcription products and 2-LTR circles was performed using qPCR. The primers and qPCR conditions were described previously (*13*). Briefly, VSV-G pseudotyped HIV-1 virions produced from HEK293T cells were quantified by HIV-1 p24 ELISA (ZeptoMetrix) and treated with 60 U/ml DNase I (Ambion) to remove the residual plasmid. MT4 cells (2×10^5^) were infected with VSV-G pseudotyped HIV-1 virions (500 ng of p24). The cells were collected at 12 h post-infection (hpi). Genomic DNA was isolated by using DNeasy Blood&Tissue kit (Qiagen). All the samples were normalized to GAPDH and 2–ΔΔCt method was used for relative quantification analysis.

To examine effects of aphidicolin on infection of WT and M66I viruses, MT4 cells were treated with 1 μg/ml aphidicolin 16 h before infection. 5 × 10^4^ cells/well (aphidicolin pretreated or naïve control) were seeded in a 24-well dish and infected with 10 ng of p24 WT and 100 ng of p24 M66I luciferase-encoding pseudotyped HIV-1 viruses. Supernatants were removed and cells were washed with PBS once after 1 hpi. Fresh complete medium with and without aphidicolin were added and the cells were cultured further for 48 h. The cells were lysed with the reporter lysis buffer (Cat.# E1531, Promega) and centrifuged to remove the cell debris. Luciferase activity in cellular extracts was determined using Luciferase Assay System (Cat.# E1531, Promega). MLV was used as a control for aphidicolin treatment.

### TEM of virions

HEK293T cells, obtained from America Type Culture Collection (cat. # CRL-3216), were monitored for mycoplasma using a commercial kit (MycoAlert PLUS, Lonza). Experiments were conducted using mycoplasma-free cells, which were propagated in Dulbecco’s modified Eagle’s medium containing 10% fetal bovine serum, 100 IU/ml penicillin, and 100 π g/ml streptomycin (DMEM) at 37°C in the presence of 5% CO_2_. Viruses were generated by transfecting cells (10^7^ plated in 15-cm plates the previous day) with 30 µg pNL4-3 plasmid DNA (WT or M66I) using PolyJet DNA transfection reagent (SignaGen Laboratories) according to the manufacturer’s protocol. Two days post-transfection, virus-containing cell supernatants were clarified by centrifugation at 800xg for 15 min and subsequently filtered by gravity flow through 0.45 µm filters. CA levels were assessed by p24 ELISA using a commercial kit (Advanced Bioscience Laboratories, Cat. #5447). Viruses were pelleted by ultracentrifugation using a Beckman SW32-Ti rotor at 26,000 rpm for 2.5 h at 4°C. Virus pellets resuspended in 1 mL fixative solution (2.5% glutaraldehyde, 1.25% paraformaldehyde, 0.03% picric acid, 0.1 M sodium cacodylate, pH 7.4) were incubated at 4 °C overnight. The preparation and sectioning of fixed virus pellets was performed at the Harvard Medical School Electron Microscopy Core Facility as previously described (*52*). Sections (50 nm) were imaged using a JEOL 1200EX transmission electron microscope operated at 80 kV. All images used for quantitative analysis were taken at 30,000-fold magnification. Images were uploaded to ImageJ and analyzed using the Cell Counter plugin, which allows for numbered markers corresponding to categories to be assigned to individual virions. Only virions that were within the plane of focus with an electron-dense outer membrane were considered for quantification. All virions were assigned one of seven morphological categories: (1) mature, (2) immature, (3) empty, (4) rod-shaped core, (5) multiple cores, (6) multiple densities (when at least one of the densities could not be attributed to a core structure), and (7) empty core. The characterization of mature and immature virions was based on previously published descriptions of these phenotypes (*52*). All other categories were established based on qualitative observations made prior to image quantification. A cumulative category, labeled "atypical" and comprising of categories 4-7, was assessed as an all-encompassing category for irregular phenotypes. Examples and descriptions of each morphological category are provided in Fig. 4A. Within each experiment we counted minimally 200 virions per condition. The number of virions per category was summed and normalized as a percentage of total virions within that set. Unpaired Student’s t-tests were performed in Prism Version 9.0.1 to assess differences in the mean frequencies for each morphological category as compared to the WT.

### Cells and plasmids

pNL4-3, a lab-adapted, subtype B infectious molecular clone (*53*), was used for transfection of SupT1 and HEK293T cells. The pNL4-3/KFS (env-) molecular clone (*54*) was used as a non-replicating control. SupT1 and HEK293T cell lines were acquired from the American Type Culture Collection (ATCC). SupT1 cells were cultured in Roswell Park Memorial Institute (RPMI) medium (Corning) supplemented with 10% fetal bovine serum (GenClone), 2 mM L-glutamine, penicillin (100 U/mL; Gibco) and streptomycin (100μg/mL; Gibco). HEK293T cells were cultured in Dulbecco’s Modified Eagle Medium (DMEM) supplemented as above.

### Forced evolution experiments

Transfection of SupT1 cells (Fig. S2) with pNL4-3/KFS (WT or M66I) was carried out using 0.7 mg/ml DEAE dextran as described previously (*55*). Cultures were split (1:2) and supernatant was collected every 2-4 days for up to three months. Replication kinetics were determined by quantifying HIV-1 reverse transcriptase activity present in supernatant samples using a previously described ^32^P-based assay (*56*). Cells and supernatant were collected at the peak of replication and the HIV-1 GagPol coding region was amplified from genomic DNA for sequencing. Sequencing results were evaluated using SnapGene.

WT or M66I viruses were produced in HEK293T cells via transfection of respective NL4-3 molecular clones. Linear polyethyleneimine (PEI) was used as a transfection reagent and virus was quantified by the above-mentioned ^32^P-based RT assay. SupT1 cells were infected for two hours at 37 °C using 0.1 RT counts per minute (cpm)/μL per cell. Replication kinetics and sequencing were performed as described above. LEN was acquired from MedChemExpress, resuspended in 100 % DMSO, and stored at -80 °C until use.

### Fluorescent virus production

Fluorescently tagged viruses were produced in HEK293T cells by transient transfection of 2 µg pHIV-(WT or M66I) alongside plasmids encoding for the pVpr-INmNeonGreen (0.8 µg) and VSV-G envelope glycoproteins (0.5 µg), using JetPrime transfection reagent in a single well of a 6-well plate, and as per manufacturers protocols. Where indicated the plasmid encoding the capsid marker pCyclophilinA-DsRed (CDR, 0.8 µg) was co-transfected to add a secondary capsid label to the virus for uncoating studies (Fig. 5). The virus supernatants were collected by brief centrifugation and by filtration through a 0.45 µm filter to remove cellular debris. All viral preps were quantified by the product-enhanced reverse transcription (PERT) assay, aliquoted, and stored at -80°C until use.

### Capsid stability measurements *in vitro* and in living cells

For in vitro core stability measurements, HIV-1(WT or M66I) pseudovirus particles fluorescently labeled with the INmNG (vRNP-marker) and CDR (capsid marker) were immobilized on poly-l-lysine treated 8-well chambered slide (#C8-1.5H-N, CellVis) by incubating at 4°C for 30 min. Unbound virus particles were removed followed by 1x wash in PBS. The bound virus particles were imaged on a Leica SP8 LSCM, using a C-Apo 63x/1.4NA oil-immersion objective. Tile-scanning was employed to image 2 x 2 neighboring fields of view. Saponin (100 µg/ml) was added to permeabilize the virus membrane and initiate capsid uncoating *in vitro.* Time-lapse images were collected at 1-min intervals for 30 min to visualize the loss of CDR from INmNG-labeled cores. Cyclosporin A (CsA, 5 μM) was added at 25 min after permeabilization/onset of imaging to displace the CDR/capsid marker from permeabilized viral cores that have not disassembled during the imaging time window.

*In situ* capsid stability in living cells was determined as described previously (5). Briefly, VSV-G pseudotyped HIV-1 particles co-labeled with INmNG and CDR were used to infect TZM-bl cells (MOI 0.008). Virus was allowed to bind to 5x10^4^ TZM-bl cells adhered to an 8-well chamber slide (#C8-1.5H-N, CellVis) by spinoculation for 30 min at 1500×g, 12°C. Before virus binding, cell nuclei were stained for 10 min with 2 μg/ml of Hoechst-33342. The cells were washed twice, and virus entry was synchronously initiated by adding pre-warmed live-cell imaging buffer (Invitrogen) supplemented with 10% FBS, followed by incubation in a CO_2_ incubator maintained at 37°C. Virus-bound cells were imaged on a Leica SP8 LSCM by acquiring images from 4 neighboring fields of view, at a frequency of 15-min intervals starting from 0-2.5 hpi. CsA (25 μM) was added at 2 h, and imaging was continued for an additional 30 min to discriminate between post-fusion cores in the cytoplasm that lost CDR in response to CsA addition, from endosomal resident intact virions that do not release this marker. The number of INmNG and CDR puncta in pre-and post-CsA treatment images was determined using the spot detector plugin in ICY. The number of stable cores residing in the cytoplasm was determined by single particle-based detection of CDR loss from INmNG puncta upon CsA addition. Images were processed offline for further analysis (see below).

### Live-cell imaging to measure capsid stability, docking at the nuclear envelope, and nuclear import

Single HIV-1 infection in live cells was visualized as previously described (*35, 57*). In brief, 5×10^5^ TZM-bl cells that stably express the emiRFP670-laminB1 nuclear envelope marker were plated in a 35 mm 1.5 glass bottom dish (#D35-20-1.5-N, CellVis). Aphidicolin (Sigma Aldrich) was added to cells (10 µM final concentration) to block cell division. About 14 h later the cells were infected within either INmNG or INmNG/CDR co-labeled fluorescent HIV-1 pseudo particles (WT or M66I) at a MOI 0.2 for long-term NE-interactions, nuclear import and infection studies. Virus binding onto cells was augmented through spinoculation (1500×g for 30 min, 16°C). Following spinoculation, virus entry was synchronously initiated by adding a pre-warmed complete DMEM medium containing Aphidicolin (10 µM) to samples mounted on a temperature-and CO_2_-controlled microscope stage.

3D time-lapse live cell imaging was carried out on a Leica SP8 LSCM, using a C-Apo 63x/1.4NA oil-immersion objective. For capsid stability measurements time-lapse imaging was performed starting from 0 – 2.5 hpi every 15 min, over 2x2 fields. CsA (5 μM) was added at 2 hpi, and the loss of CDR from cytoplasmic capsids was imaged for an additional 30 min. For HIV-NE interaction studies, live-cell time-lapse imaging was performed starting from 0.5 – 16 hpi every 2.5 min, by using tile-scanning of 4x4 fields of view. Imaging was performed by acquiring 9-11 Z-stacks spaced by 0.8 μm. Images were collected using the following setting: 512x512 pixels/frame, 180 nm pixel sizes, scan speed of 1.54 µs pixel dwell time. Highly attenuated 488, 561, and 633 nm laser lines were used to excite the INmNG, CDR, and emiRFP670-Lamin B1 fluorescent markers, and their respective emission was collected between 502-560 nm, 570-630 nm, and 645-700 nm using GaSP-HyD hybrid detectors. HIV-1 nuclear import (8 hpi) and infection (eGFP-expressing cells, 16 hpi) were quantified from live-cell movies. 3D-image series were processed offline using ICY image analysis software (http://icy.bioimageanalysis.org/) (*58*).

### Image analyses

#### Capsid stability measurements

For *in vitro* core-stability measurements, the number of INmNG and CDR puncta was detected, using the wavelet spot detector in ICY. The kinetics of *in vitro* CA disassembly was determined by normalizing the number of CDR puncta to that at t=0 min and subtracting the background CsA-resistant CDR puncta at t=30 min (CsA was added at 25 min), presumably corresponding to immature viruses. The total number of INmNG puncta in each field of view remained constant over the 30-min imaging period. The capsid stability was determined by plotting the loss of CDR spots over time, normalized to the initial number of spots. Immature particles that retained CDR and failed to respond to CsA treatment were excluded from the analysis.

For *in situ* capsid stability measurements, the 3D-confocal z-stacks of live-cell time-lapse movies were converted to a 2D-maximum intensity projection. The number of INmNG and CDR puncta associated with the individual cells of the time series was determined using the wavelet spot detector plugin in ICY. The initial fraction of cell-associated INmNG/CDR labeled cores and the final fraction of cytoplasmic cores, which responded to 25 µM CsA treatment (*added at 2 hpi*) and had lost the CDR/capsid marker were calculated to determine the total fraction of capsids that productively entered the cell through virus-cell fusion. The fraction of CDR puncta loss (capsid stability) in the cytoplasm at each time point of the live-cell movie was normalized to the total cores that entered the cytoplasm. Endosomal cores that did not respond to CsA treatment, and did not lose CDR, were excluded from the analysis.

#### HIV-1 interactions with the NE

For analysis of HIV-1 docking at the NE, we selected >10 cells from each independent live-cell experiment that showed minimal lateral movements. The initial (0-4 h) when there is a higher density of INmNG puncta in the cytoplasm, and the latter (12-16 h) segments showing extensive cellular motion in the live-cell movies were removed from the analysis. The central 8 h segment (between 4-12 hpi) was chosen to facilitate robust, high-confidence particle tracking. Analysis was further delimited to the central z-planes (2x planes, 1.6 µm) of these nuclei, which were extracted and projected to a 2D stack. Projected 2D-images were additionally drift-corrected using Fast4DReg in Fiji (*59*) and were denoised using N2V (*60*). The laminB1 fluorescence signals in the processed images were used to mask the pixels occupied by the NE to create a region of interest (ROI). The HIV-1 INmNG puncta detected within this NE mask was tracked in an automated fashion in the ICY bio-image analysis single particle tracking suite.

Tracks that correspond to 3 or more frames (≥7.5 min) were considered as NE interactions. Each time point in a track was used as a center to determine the number of consecutive frames that a single HIV-1 particle remained localized within a 2-pixel (360 nm) radius for 3 or more frames. These tracks were deemed as stable interactions (or) docking at the NE. The 360 nm radius was used taking into consideration the lateral diffraction-limited confocal resolution (∼240 nm) and the pixel sizes (180 nm) in our time-lapse movies. These stable interaction events, as a fraction of total NE interactions obtained in each nucleus over the 8 h time window were analyzed and plotted. As controls, the interactions between non-specific interactions between bald endosome-trapped HIV-1 (no-VSV) infections and the NE were analyzed. Of the 15.8% of non-specific interactions detected by this method, only 5.1% docked at the NE for longer than 7.5 min, which was >5-fold lower than interactions between capsids (WT and M66I). HIV-1 nuclear import in fixed time-point 3D z-stack images was analyzed using an in-house script in the ICY protocols’ module as described in (*35*). Briefly, the inner nucleus volume was segmented and the number of HIV-1 INmNG puncta inside the nucleus was detected using the HK-means (for segmentation), and spot-detector (for fHIV-1 puncta detection) plugins in ICY.

### Sequencing of integration sites

Integration site libraries were prepared by performing ligation-mediated PCR (LM-PCR) as described previously with minor modifications (*13, 61*). Briefly, HEK293T cells were seeded (2x10^5^ cells/well) in a 6-well plate 24 h prior to infection. On day 1, the cell culture medium was replaced with a fresh DMEM medium containing polybrene (8 µg/ml). Cultured cells were infected with VSV-G pseudotyped HIV-1 luciferase virus (pNL4-3E-R+luc, WT or M66I, 2000-6000 ng of p24) and incubated for 2 h at 37°C, 5% CO_2_. Then, the virus-containing medium was removed and replaced with a fresh DMEM medium. The cells were incubated at 37 °C, 5% CO_2_ for 48 h. Then cells were harvested, and 30% of cells were used to measure luciferase activity with Luciferase Assay System (Promega). The remaining cells were cultured for 5 days in a 150 mm cell culture dish. After 5 days cells were harvested, genomic DNA was extracted using QIAamp DNA Mini Kit (Qiagen) and 10 µg genomic DNA was digested with MseI and BglII restriction enzymes. Digested DNA was purified by QIAquick PCR Purification Kit for PCR Cleanup kit (Qiagen) and used to set up 4-independent ligation reactions to a double-stranded asymmetrical linker with a 5’-TA overhang.

Purified ligated DNA was used to amplify virus-host junctions with linker and HIV-1 U5 specific LTR primers in a semi-nested PCR reaction, as described (*62, 63*). Purified PCR amplicons were sequenced on the Illumina platform at Genewiz. FASTQ raw reads were trimmed for HIV-1 U5 and linker sequences, the trimmed reads were aligned to the hg19 human genome, filtered, and unique integration sites were selected. Integration sites (%) were calculated for genomic features such as genes, transcription start sites, CpG, LADs and SPADs. Random integration control (RIC) contains *in silico* generated random sites from the hg19 genome and were previously published (*64*). P-values were calculated in pairwise comparisons for genes, TSS, LADs, SPADs, and CpG by Fisher test and by Wilcoxon test for gene-density (*63*).

## Data availability

The coordinates of the crystal structures of CA(M66I)_Hex_ and CA(M66I)_Hex_+IP6+CPSF6_313-327_ have been deposited at the Protein Data Bank (PDB) under accession numbers 7RAO and 8GDV. Raw FASTQ sequences generated for integration sites mapping in this study are available in the National Center for Biotechnology Sequence Read Archive (NCBI SRA) under accession number SRR31330709. All data needed to evaluate the conclusions in the paper are present in the paper and Supplementary Materials. All materials will be made available upon request.

## Acknowledgments

We are grateful to Garry Morgan and Anza Darehshouri for the technical suggestions for the electron micrographs. Electron micrographs were collected at Boulder Electron Microscopy Services at the University of Colorado, Electron Microscopy Core Facility at the University of Colorado Anschutz Medical Campus, and Harvard Medical School Electron Microscopy Core Facility. We thank Jay Nix at ALS Beamline 4.2.2 for acquiring data for the crystal structure.

## Author contributions

Conceptualization: SWH, EOF, ANE, ACF, MK. Investigation: LB, ASA, SMB, GW, JRAM, SPS, ABK, RR, PKS, JG, LS, RH, SWH, MT, BN. Supervision: SWH, EOF, ANE, ACF, MK. Writing, review and editing: all authors.

## Declaration of interests

The authors declare no competing interests.

## Financial Disclosure

This work was supported by NIH grants R01 AI157802 and R01 AI162665 (MK), U54 AI170855 (MK and ACF), R01 AI181627 and R21 AI174879 (ACF), R01 AI052014 (ANE), U54 AI170791 (ANE), T32 AI150547 (SWH), K99 AI174891 (ABK). Research in the Freed laboratory is supported by the Intramural Research Program of the Center for Cancer Research, National Cancer Institute, National Institutes of Health.

